# Characterization of nucleus accumbens dopamine dynamics during positive and negative operant reinforcement in rats

**DOI:** 10.64898/2026.07.24.740421

**Authors:** Jonathan J Chow, Emma M Pilz, Huiling Wang, Kauê M Costa, Geoff Schoenbaum, Yavin Shaham

## Abstract

We previously reported, using *in-vivo* fiber photometry, that operant responding reinforced by access to a peer (social self-administration) is associated with phasic dopamine increases in nucleus accumbens (NAc) core following lever insertion (reward-availability cue) and gradual increases preceding lever-pressing. Here, we sought to replicate these findings and determine whether dopamine signals (1) generalize to responding for high-carbohydrate palatable food, (2) show opposite patterns during negative reinforcement (shock avoidance/escape), and (3) depend on whether reinforcers are experienced alone or together.

We trained rats (n=11; 6 females) to lever-press for access to a same-sex peer (15 s/trial) and palatable food (45-mg pellet/trial), followed by shock avoidance/escape (0.18–0.26 mA). After training, we expressed the dopamine sensor GRAB-DA2m and implanted optic fibers into NAc core. We measured dopamine activity during sessions with either one- or three-reinforcers.

During social self-administration, dopamine activity showed phasic increases following lever insertion and gradual increases preceding lever-pressing; responses were moderately greater during sessions with all three reinforcers. Palatable food self-administration showed a similar pattern, but responses were approximately twofold greater during single-reinforcer sessions. During shock avoidance/escape, dopamine activity showed phasic decreases at warning onset, lever insertion, and shock onset; responses were also greater during single-reinforcer sessions.

Results suggest that NAc core dopamine signaling distinguishes positive from negative reinforcement and is modulated by reinforcer availability. Compared with single-reinforcer sessions, dopamine responses during food self-administration and shock avoidance/escape were reduced during sessions with all three reinforcers, whereas responses during social self-administration modestly increased.

**Significance statement:** Nucleus accumbens (NAc) dopamine is critical for processing the valence of positive reinforcers, whereas its role in negative reinforcement is less well understood. Here, we extended our prior work on operant social self-administration to determine whether these findings generalize to food reinforcement, show the opposite pattern during operant negative reinforcement (shock avoidance/escape), and depend on whether reinforcers are experienced alone or together. Results suggest that NAc core dopamine activity differentiates positive and negative reinforcement and is modulated by the presence of other available reinforcers. Specifically, dopamine responses during food self-administration and shock avoidance/escape were reduced when all three reinforcers were available together compared with when each reinforcer was experienced alone, whereas responses during social self-administration showed the opposite pattern.

## Introduction

The robust inhibitory effects of operant social interaction on drug reward and relapse (Venniro et al., 2018; Venniro et al., 2019; Venniro et al., 2021) motivated us to investigate the neurobehavioral mechanisms underlying operant responding for social interaction (social self-administration) (Chow et al., 2022; Chow et al., 2024b). We found that the duration of peer access had little effect on social self-administration, whereas operant responding was higher for a familiar than for a novel peer and decreased as the fixed-ratio requirement for access to the social reward increased. We also showed that rats strongly preferred high-carbohydrate palatable food over social interaction and were willing to complete higher fixed-ratio requirements to obtain the food reward. These findings indicate that social interaction functions as an operant positive reinforcer, but its reinforcing efficacy is lower than that of high-carbohydrate palatable food.

It is well established that striatal dopamine plays a critical role in the rewarding effects of both drug and non-drug reinforcers (Kelley and Berridge, 2002; Wise, 2004), including social interaction (Gunaydin et al., 2014; Vanderschuren et al., 2016; Dai et al., 2022; Solié et al., 2022). In a recent study, we characterized dopamine signaling in the nucleus accumbens (NAc) core and dorsomedial striatum (DMS) using *in vivo* fiber photometry during social self-administration (Chow et al., 2024b). We found that peer sex influenced operant responding in male but not female rats, with males responding more for opposite-sex than same-sex peers, whereas the estrous cycle had no effect on responding. We also found that phasic dopamine responses were larger following lever insertion, the cue signaling social reward availability, than following lever-pressing. These cue-evoked responses were greater in males than females. In addition, dopamine activity gradually increased during the 1 s preceding lever-pressing in the NAc core, but not the DMS. Together, these findings indicate that dopamine signaling is engaged by cues predicting social reward and by the operant response used to obtain it in a sex- and region-specific manner (Chow et al., 2024b).

Many studies have reported phasic increases in NAc dopamine during operant responding for both drug and non-drug positive reinforcers (Phillips et al., 2003; Roitman et al., 2004; Mohebi et al., 2019; Pribiag et al., 2021). Additionally, three studies reported similar findings in rats trained to lever press to avoid shock using an operant negative-reinforcement procedure (Oleson et al., 2012; Gentry et al., 2016; Wenzel et al., 2018). Therefore, the goal of the present study was to extend our previous fiber photometry studies of social self-administration by characterizing NAc core dopamine signaling during operant responding for another positive reinforcer (high-carbohydrate palatable food) and a negative reinforcer (shock avoidance/escape).

Here, we sought to replicate our previous findings on NAc core dopamine signaling during social self-administration and address three questions. First, we asked whether dopamine signaling around lever insertion and lever-pressing generalizes to operant responding for high-carbohydrate palatable food and whether these signals are larger than those evoked by social interaction, which rats value less than this palatable food (Chow et al., 2022). Second, we tested whether operant negative reinforcement through shock avoidance/escape produces a pattern of dopamine signaling opposite to that observed during positive reinforcement. Third, we asked whether reinforcer availability modulates dopamine signaling by comparing responses to social interaction, high-carbohydrate palatable food, and shock when each reinforcer was experienced alone or in sessions containing all three reinforcers. Overall, we tested whether NAc core dopamine signaling encodes reinforcer valence and whether reinforcer availability modulates these signals across positive and negative reinforcement.

## Materials and Methods

### Subjects

We used 5 male and 6 female Sprague Dawley rats with accompanying sex-matched peers (n=11; 6 females). Rats arrived pair-housed at an estimated age of ∼7 to 8 weeks PND (body weight at the time of arrival: males, 201–225 g; females, 176–200 g; Charles River). We habituated them to the animal facility for ∼1 week under a reverse 12:12 h light/dark cycle (lights off at 8 AM) with free access to food (Teklad Rat Diet, Envigo) and water. Afterwards, we single-housed the rats. We then handled the rats daily the week prior to the start of the experiment. We performed all experiment in accordance with the NIH Guide for the Care and Use of Laboratory Animals (8th edition), under protocols approved by the NIDA IRP Animal Care and Use Committee.

### Apparatus

We conducted the experiment in operant chambers attached to a partner chamber (ENV-00CT-SOCIAL; Med Associates) controlled by Med-PC IV. A guillotine door (ENV-10B2-SOC) and a perforated metal divider separated the two chambers, preventing the rats from crossing into the other chamber while allowing for full face-to-face and forepaw contact. We outfitted the chambers with recessed pellet receptacles (ENV-200R1M) containing a head-entry detector (ENV-254-CB) that was attached to a pellet dispenser (ENV-203 M-45). In half of the chambers, the food receptacle combination was placed left of the guillotine door on the same panel. Under this configuration, we outfitted the opposite panel with three retractable levers (ENV-112CM); above the right- and left-levers were white cue-lights (ENV-221 M) and above the middle lever, was a red houselight. At the very top of panel above the right-lever and cue-light was a tone generator (ENV-223 AM; 2.9 kHz).

In the other half of the chambers, the food receptacle combination was placed opposite to the guillotine door, in the middle-section. Under this configuration, we placed a red houselight above the food receptacle and we outfitted the right- and left-side of the food receptacle with levers with corresponding white-cue lights above each of them; at the very top of the panel above the right-lever was a tone generator. Next to the guillotine was a third retractable lever. All grid floors of the main chamber were wired to a shock scrambler (ENV-414). Note: directions (e.g., left, right) are from the perspective of a subject directly facing said panels.

### Behavioral training

#### Shaping

On the first day of the experiment, we first trained the rats for door shaping. We assigned each experimental rat with a same-sex peer that was different from arrival; this assigned-rat would serve as the peer for the rest of the experiment. The door shaping procedure began with a 15-min acclimation phase followed by a phase in which the guillotine door separating the experimental and peer rat was lifted for 15 s, allowing for contact through the cutouts in the metal divider. After 15 s, the guillotine door closed and a 60 s intertrial interval (ITI) separated the next trial; this repeated for a total of 15 trials. After the last trial, rats remained in the chambers for 5-min to continue acclimation. After 5-min, we removed the peer rats from the partner chambers and initiated food magazine training. The magazine shaping procedure began with a 15-min acclimation phase followed by a phase in which a single 45-mg palatable pellet (TestDiet, 5TUL-1,811,155, 12.7% fat, 66.7% carbohydrate, and 20.6% protein) would be dropped into the food receptacle every 75 s for a total of 15 pellets. Rats remained in the chambers for an additional 5 min after the last pellet delivery to continue acclimation.

#### Social and food self-administration

After door- and magazine-shaping, we trained the rats for social and food self-administration on a fixed-ratio (FR) 1 schedule of reinforcement in separate back-to-back sessions (Figure 1A). Each session began with a 3-min dark period. Next, a single-lever extended signaling the start of a trial. The lever assigned to social reinforcement was opposite to the guillotine door and the lever assigned to food reinforcement was opposite to the food receptacle. Rats in chambers with the food receptacle and social door on the same panel, lever-pressed on either the middle- or left-lever for social interaction and food (counterbalanced). Rats in chambers with food receptacle and social door on opposite panels, lever-pressed on the lever next to the guillotine door for food and either the left- or right-lever for social interaction (counterbalanced). During each trial, completion of the FR1 requirement resulted in the retraction of the lever and delivery of the reinforcer after a 0.5-s delay. For social interaction, the guillotine door would open for 15 s; for food, a single 45-mg pellet would be dispensed. Social trials were separated by a 20 s ITI whereas food trials were separated by a 35 s ITI. Sessions lasted for 45 min or ended when 45 reinforcers were earned. We trained the rats for social interaction first, after which we removed the peer rats, and initiated food training. We trained the rats for 10 d.

**Figure 1.**
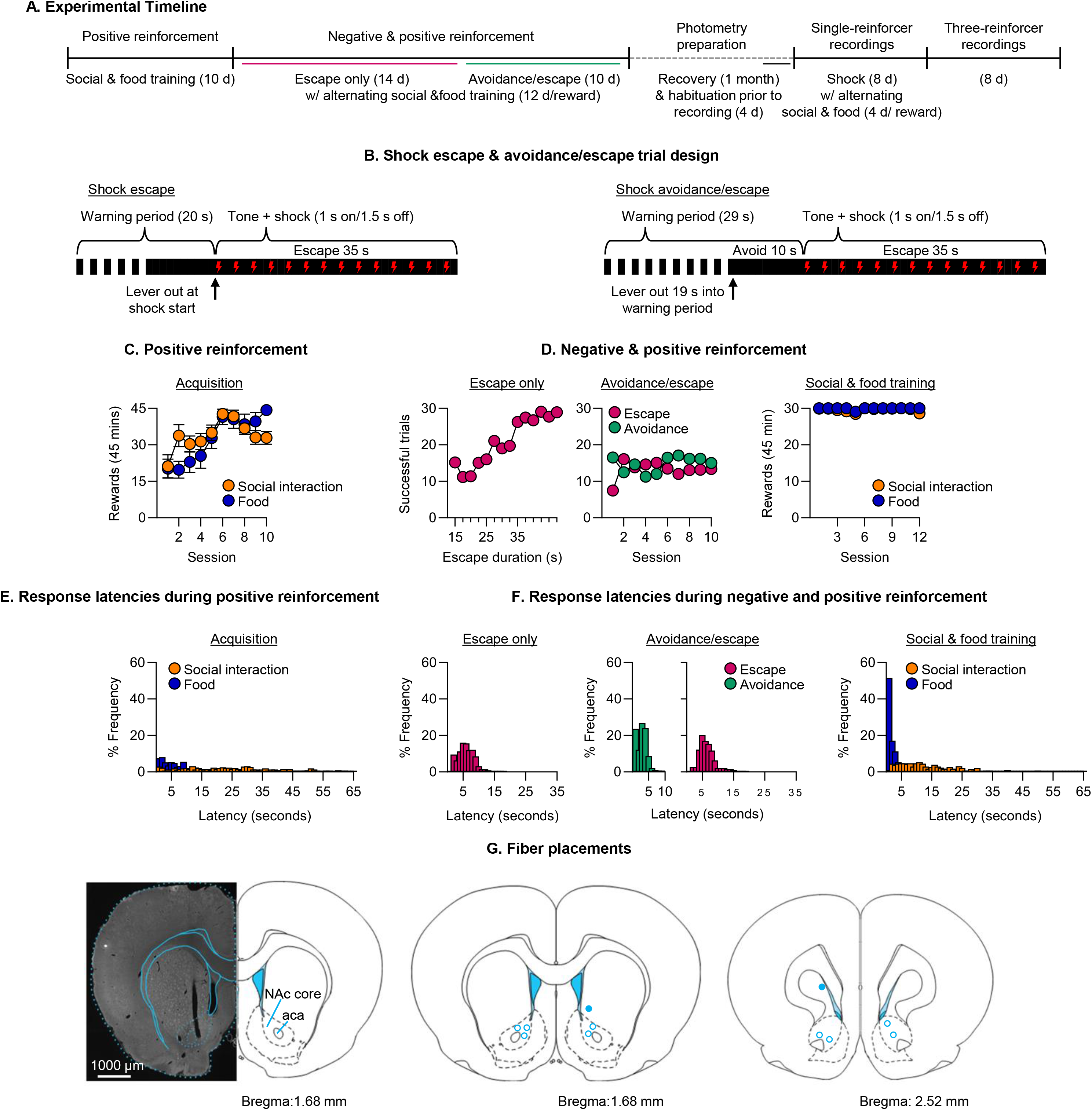
Timeline, behavioral training, and fiber placement. Operant responding for food and social interaction and for shock escape and avoidance. **(A)** Experimental timeline. **(B)** Schematic of the shock escape trials and shock avoidance/escape trials. **(C)** Number of food and social interaction rewards earned during acquisition. **(D)** Number of successful escape trials during training, number of successful avoidance and escape trials during shock blocks, and number of food and social interaction rewards earned during the 12 days preceding photometry surgery, demonstrating stable responding. **(E)** Latency to lever-press following lever extension during food and social interaction blocks. **(F)** Latency to lever-press following lever extension during escape-only, avoidance/escape, and food/social blocks. **(G)** Representative brain section and optic fiber placements in the nucleus accumbens core (NAc core). n = 11 (6 females).

#### Initial shock lever training and shock assessment

During the last 4 d of social and food self-administration (days 7 to 10 of social and food training), we introduced the third lever (deemed shock lever; whichever lever the rats have not encountered yet) as part of initial shock training (Chow et al., 2024a). The shock lever would extend randomly (5 times) during the first 25 min of each social and food training session. Completion of an FR1 would retract the shock lever and allow rats to resume social and food self-administration. If a lever-press was not made within 4 min, the shock lever would retract, and rats could resume social and food self-administration. In addition, during the last day of social and food self-administration training, we assessed each rats’ shock sensitivity. After placing the experimental rats into their respective operant chambers, we set the associated shock scramblers for each respective chambers to 0.1 mA and turned them on for ∼ 3 s. During the shock assessment, we observed the rats for backwards-scurrying and/or lifting of the front paws. If neither of these behaviors were observed, we increased the shock intensity by 0.01 mA until the target behavior was observed. The initial shock intensity for the rats ranged from 0.14 to 0.16 mA. After which, we placed the peers into the partner side and initiated social self-administration training.

#### Shock escape training

Next we trained rats for shock escape based on prior methods from the lab (Chow et al., 2024a). Each session began with a 3-min dark period. Next, a warning period would start, which signaled the start of a trial. The warning period consisted of a tone a beeping (1-s on/1-s off) over 10 s, then transitioned into a solid tone for another 10 s. At the end of the 20-s warning period, in continuing presence of the tone, the shock lever would extend into the chamber and intermittent foot-shock (1-s on/1.5-s off; at the previously determined intensity) began. During this period, we allowed the rats 15 s to 35 s (4 d at 15 s, 4 d at 25 s, and 6 d at 35 s) to complete the FR1 lever-press requirement to terminate the shock and tone (Figure 1B, left). After which, the trial would enter a 20-s ITI plus whatever duration remained of the shock period.

If the rats did not emit a response during the allotted duration, the shock lever would retract, and the shock and tone would terminate and enter a 20-s ITI. This cycle repeated for a total of 30 trials. During initial shock escape training we placed large acrylic boxes into the operant chambers to reduce the area in which the rats could move and to force them to be closer to the shock-paired lever. This box was removed once rats started achieving > 12 escapes/session. We also increased the shock intensity (0.01 to 0.03 mA/session) if a rat failed to achieve an average of > 5 successful escapes over the previous two days of training. If increasing the shock intensity resulted in vocalization or jumping responses, we decreased the footshock intensity or left it as is; the shock intensity at the end of shock escape training ranged from 0.16 to 0.24 m. After each shock escape session, we continued to train rats for social and food self-administration, as described above, alternating days between the two reinforcers (maxed at 30 reinforcers/ 45-min session). We placed the peer rats into the partner chambers only after the shock escape session ended.

#### Shock avoidance/escape training

After shock escape training, we advanced all the rats to an active avoidance and escape procedure (Chow et al., 2024a). Each session began with a 3-min dark period. Next, a warning period would start, which signaled the start of a trial. The warning period consisted of a tone a beeping (1-s on/1-s off) over 19 s, then transitioned into a solid tone for another 10 s which coincided with the extension of the shock lever (Figure 1B, right). At the end of the 29-s warning period, in continuing presence of the tone, intermittent foot-shock (1-s on/1.5-s off) began and lasted up to 35 s. Completion of the FR1 lever-press requirement during the 10 s prior to shock onset counted as avoidance response and resulted in termination of the tone and avoided shock altogether.

Lever-press after shock onset counted as an escape. After successful avoidance or escape, the trials entered a 20-s ITI plus whatever remaining time was left for the trial. Failure to lever-press resulted in the shock and tone terminating and entered a 20-s ITI. This cycle repeated for a total of 30 trials. We trained rats on shock avoidance/escape for 10 sessions. If rats showed greater than 5 omissions (i.e., failure to avoid or escape) per session, we would increase the shock intensity (0.01 to 0.03 mA/session). The shock intensity at the end of shock avoidance/escape training ranged from 0.18 mA to 0.26 mA. After each shock avoidance/escape session, we continued to train the rats for social and food self-administration, as described above, alternating days between the two reinforcers (maxed at 30 reinforcers/ 45-min session). We placed the peer rats were placed into the partner chambers only after the shock avoidance/escape session ended.

#### Surgery

After training, we performed bilateral intracranial surgery for *in vivo* fiber photometry on the experimental rats. We anaesthetized the rats with isoflurane gas (5% induction, 2-3% maintenance). We bilaterally injected a 1:1 mixture of GRAB_DA_ (pAAV-hsyn-GRAB_DA2m from Addgene #140553) and GRAB_r5HT1.0_ (pAAV-hSyn-GRAB_r5-HT1.0 from Addgene #208719;) into the NAc core (AP +1.7 mm, ML ± 1.7 mm, and DV −6.4/-6.3, from brain surface) and the NAc shell (AP +1.68 mm, ML ± 0.75 mm, and DV −6.7/-6.6, from brain surface). We counterbalanced the injections and implants into the NAc core and NAc shell across the hemispheres.

We injected 0.7 µL of the mixture of GRAB_DA_ + GRAB_r5HT1.0_ virus at each site (NAc core DV −6.4 & −6.3; NAc shell DV −6.7/-6.6) at 0.2 μl/min using an infusion pump. We then implanted optic fibers (10 mm length, 200 μm diameter, 1.25 mm ferrule; Neurophotometrics) in each corresponding region located within the viral infusions (NAc core DV −6.34; NAc shell DV −6.64) and secured them with modified metal protective headcaps (Thorlabs) to the skull with dental cement and jeweler screws. We injected the rats with ketoprofen (2.5 mg/kg, s.c., Covetrus) after surgery and the following 3 d to relieve pain and inflammation; we also injected the rats for a week following surgery with gentamicin (5 mg/kg, s.c., APP Pharmaceuticals) to prevent infection.

#### Photometry recording

After ∼4 weeks of recovery, we habituated the rats to the photometry tether system (without connecting the fibers). We connected the rats to the tether system and trained them on a shorter version of social and food self-administration (5-min dark period followed by 15 trials), alternating between the two reinforcers over 4 days. After this period, we began formal recording. Note: We counterbalanced rats to different levers for each reinforcer, which, combined with the additional counterbalancing of the optic fiber implant locations, was designed to avoid any lateralization effects of the recorded signals.

#### Single-reinforcer recordings

We first tested rats under a single-reinforcer design. We used the same behavioral procedures for shock avoidance/escape, social self-administration, and food self-administration described above (30 trials per session; Behavioral Training). At the start of each recording session, we connected the implanted optic fibers to the photometry patch cord and initiated photometry acquisition. We first recorded a shock avoidance/escape session. We then stopped photometry acquisition and, after a ∼15 to 20 min break, initiated recordings for a social or food self-administration session, alternating the session type across days. Before each social self-administration session, we placed the peer rat into the partner chamber during the intersession interval. At the end of each session, we disconnected the rat and returned all rats to their home cages. We recorded eight shock avoidance/escape sessions and four sessions each of social and food self-administration.

#### Three-reinforcer recordings

We next tested rats under a multiple-reinforcer design in which all three reinforcers were available within a single session. At the start of each recording session, we connected the implanted optic fibers to the photometry patch cord, placed the peer rat into the partner chamber, initiated photometry acquisition, and started the behavioral program. The program presented shock avoidance/escape, social, and food trials in pseudorandom order (no more than five consecutive trials of the same reinforcer). We used the same trial structure for each reinforcer described above. We recorded eight sessions under this design, with each session including 20 trials of each reinforcer. At the end of each session, we disconnected the rat and returned all animals to their home cages.

#### Recording

We recorded dopamine and serotonin signals using a FP3002 system (Neurophotometrics) which was operated by Bonsai open-source program (Lopes et al., 2015). We used a branching fiber-optic patch cord (200 μm diameter, 0.37 NA, Doric Lenses) that allowed us to connect both implanted-fibers in the rats simultaneously (up to 4 rats/patch cord). We connected optic fibers together via shrink-wrapped protected brass sleeves (Thorlabs) and secured them using custom metal headcaps (Thorlabs) attached to a custom-made swivel-tether system that allowed for relatively free movement of rats. We pulsed a 560 nm (active signal; serotonin), 470 nm (active signal; dopamine), and 415 nm (isobestic reference) light at 100 Hz (33 Hz acquisition rate for each channel for recordings). At the end of the recording, we anesthetized the rats, perfused them, and extracted the brains. We sectioned brains coronally at 40 µm using a cryostat and verified fiber placements by microscopy. Of the samples collected, fiber placements were confirmed in the NAc core in 9 of 11 rats (5 females) and in the NAc shell in 3 of 11 rats. Due to the limited number of successful NAc shell placements, we report only the NAc core data.

#### Data processing

We processed the signals using custom scripts in MATLAB (MathWorks) (Chow et al., 2024b) to characterize dopamine transients during operant responding for social interaction, palatable food, and shock escape avoidance. We first applied a fourth-order median filter to the raw fluorescence signals from the 470 (active signal dopamine) and 415 (isosbestic reference) channels. Next, we performed a second-order polynomial fit to the reference channel data, which was then subtracted from the active channel to correct for dopamine-independent variations in fluorescence. We z-scored the resulting signal for each trial, using a 2.5-s window which was moved to 5 s prior to the start of each trial onset (e.g., warning tone for shock; lever extension of social/food) as a baseline. <u>Note:</u> We also analyzed serotonin (560 nm) signals by first filtering and detrending each session’s recordings, followed by z-scoring the resulting signal for each trial (Costa et al., 2025). However, we do not report these data in the present study because the serotonin measurements require additional validation before meaningful conclusions can be drawn.

#### Statistical analysis

We analyzed the data using linear mixed-effects (LME) models in JMP (Version 16) with α set to 0.05. In all behavioral analyses, we used subjects (the rats) as a random factor. We analyzed the neural data using the same statistical approach as in our previous paper (Chow et al., 2024b) with trials (the trials across all subjects for all sessions) as a random factor. When appropriate, we followed significant main effects and interactions with Tukey’s post-hoc tests. We included both male and female rats in accordance with NIH guidelines for considering sex as a biological variable. However, we did not include sex as a factor in the statistical analyses because the sample sizes within each sex were too small to support meaningful comparisons (single-reinforcer sessions: 5 females and 4 males; three-reinforcer sessions: 4 females and 4 males).

## Results

### Behavioral data

#### Acquisition of social and food self-administration

Rats learned to lever-press for social interaction and food (Figure 1C). We analyzed the data together using session (continuous; 1 to 10) and reinforcer type (nominal; social, food) as within-subjects factors. The analysis showed significant effects of session [F(1,10) = 32.7, p = 0.0002] and session × reinforcer type interaction [F(1,10) = 9.7, p = 0.011], due to higher responding for social interaction during the initial sessions and lower responding during the later sessions.

During training, the latency to lever-press was generally shorter for the palatable food than for social interaction (Figure 1E). A two-sample Kolmogorov-Smirnov test comparing the distributions of response latencies for palatable food (median = 2.62 s) and social interaction (median = 12.05 s) showed a significant difference, with the food latency distribution shifted toward shorter response latencies than the social interaction distribution (D⁺ = 0.27, *p* < 0.0001).

#### Acquisition of shock escape and shock avoidance/escape with continued social and food training

Rats learned to escape shock and avoid shock (Figure 1D, left & middle). We analyzed the data using session (continuous; 1 to 14) as within-subjects factor. The analysis showed a significant effect of session for shock escape [F(1,10) = 47.9, *p* < 0.0001]. Rats continued to lever-press to escape shock and acquired shock avoidance, performing at similar levels as escape. We analyzed the data using session (continuous; 1 to 10) and successful trial type (nominal; escape vs. avoid) as within-subjects factors. The analysis showed a significant effect of session [F(1,10) = 5.5, *p* = 0.04]. The median latency for shock escape during acquisition was 4.57 s. The median latencies to lever-press for shock avoidance or escape during subsequent training were 2.37 s and 4.49 s, respectively.

The rats also continued to lever-press for social and food at similar levels, earning near maximum reinforcers (30; estimated marginal means food = 29.9, social = 29.6), throughout shock escape and shock avoidance/escape training (Figure 1D, right). We analyzed the data using session (continuous; 1 to 12) and reinforcer type (nominal; social, food) as within-subjects factor. The analysis showed no main effects nor significant interactions of session and reinforcer type. During this phase, lever-press latencies for palatable food remained shorter than those for social interaction. A two-sample Kolmogorov-Smirnov test confirmed a significant difference between the latency distributions for food (median = 0.9 s) and social interaction (median = 2.47 s); the positive direction of the effect (D⁺ = 0.363, *p* < 0.0001) indicated that the food latency distribution was shifted toward shorter latencies than the social interaction latency distribution (Figure 1F, right).

#### Social and food self-administration during single-schedule recordings

Rats continued to lever-press for social interaction and food (Figure 2B, left). We analyzed the data together using session (continuous; 1 to 4) and reinforcer type (nominal; social, food) as within-subjects factors. The analysis showed a main effect of session [F(1,8) = 5.9, *p* = 0.042]. For latency, a two-sample Kolmogorov-Smirnov test comparing the distributions of response latencies for food (median = 0.87 s) and social interaction (median = 5.96 s) showed a significant difference, with the food latency distribution shifted toward shorter response latencies than the social interaction distribution (D⁺ = 0.464, *p* < 0.0001; Figure 2B).

**Figure 2.**
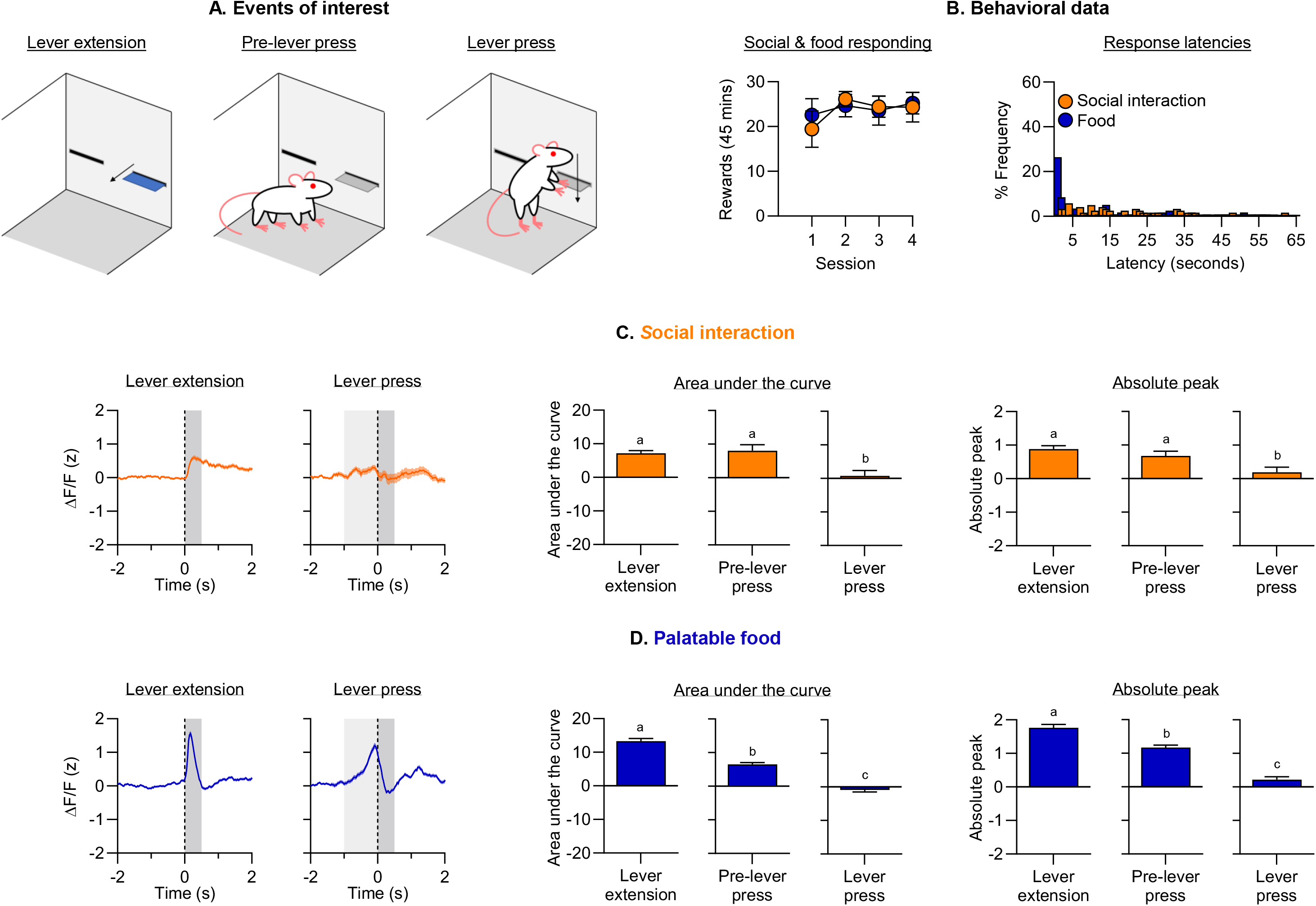
Dopamine activity in the NAc core during the one-reinforcer sessions: Food and social interaction. **(A)** Representative schematics of lever extension (left) and lever-press (right). **(B)** Number of food and social interaction rewards earned during recording sessions and latency to lever-press following lever extension during food and social interaction blocks. **(C)** Dopamine responses in the NAc core during social interaction blocks, from lever extension through the lever-press, with corresponding area under the curve (AUC) and absolute peak amplitude measures for all recorded trials. **(D)** Dopamine responses in the NAc core during food blocks, from lever extension through the lever-press, with corresponding AUC and absolute peak amplitude measures for all recorded trials. Letters indicate results of post-hoc comparisons, ranked from highest (a) to lowest; those sharing at least one letter are not significantly different. n = 9 (5 females).

#### Shock avoidance/escape during single-schedule recordings

Rats continued to avoid or escape shock (Figure 3B, top). We analyzed the data together using session (continuous; 1 to 8) and successful trial type (nominal; escape vs. avoid) as within-subjects factors. The analysis showed a significant effect of session [F(1,8) = 8.0, *p* = 0.022]. The median latencies to lever-press for shock avoidance or escape were 4.76 s and 3.12 s, respectively.

**Figure 3.**
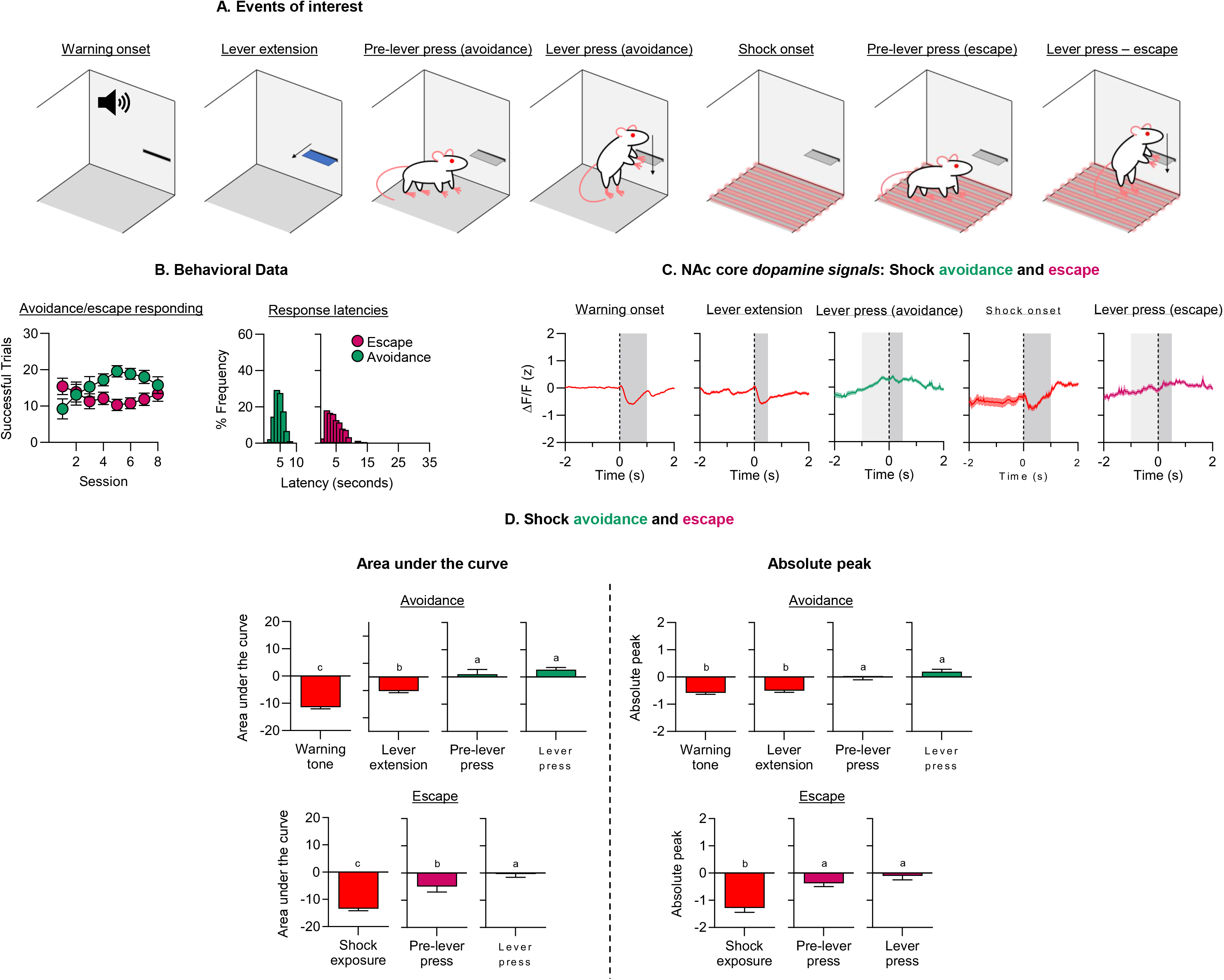
Dopamine activity in the NAc core during the one-reinforcer sessions: shock avoidance and escape. **(A)** Representative schematics, from left to right, of warning-signal onset, lever extension, the pre-press avoidance period, the avoidance response, shock onset, the pre-press escape period, and the escape response. **(B)** Number of successful avoidance and escape trials during recording sessions and latency to lever-press following lever extension during shock blocks. **(C)** Dopamine responses in the NAc core during shock blocks at, from left to right, warning-signal onset, lever extension, the avoidance response, shock onset, and the escape response. **(D)** AUC (left) and absolute peak amplitude (right) for all recorded trials at warning-signal onset, lever extension, during the pre-press avoidance period, during the avoidance response, at shock onset, during the pre-press escape period, and during the escape response. Letters indicate results of post hoc comparisons, ranked from highest (a) to lowest; those sharing at least one letter are not significantly different. n = 9 (5 females).

#### Shock avoidance/escape, social and food self-administration during multiple-schedule recordings

Rats continued to lever-press for social interaction and food (Figure 4A). We analyzed the data together using session (continuous; 1 to 8) and reinforcer type (nominal; social, food) as within-subjects factors. The analysis showed no main effects nor significant interactions of session and reinforcer type. For latency, a two-sample Kolmogorov-Smirnov test comparing the distributions of response latencies for food (median = 1.29 s) and social interaction (median = 1.75 s) showed a significant difference, with the food latency distribution shifted toward shorter response latencies than the social interaction distribution (D⁺ = 0.464, *p* < 0.0001; Figure 4A).Rats also continued to avoid or escape shock (Figure 4B). We analyzed the data together using session (continuous; 1 to 8) and successful trial type (nominal; escape vs. avoid) as within-subjects factors. The analysis showed no main effects nor significant interactions of session and successful trial type. The median latencies to lever-press for shock avoidance or escape were 3.06 s and 3.21 s, respectively.

**Figure 4.**
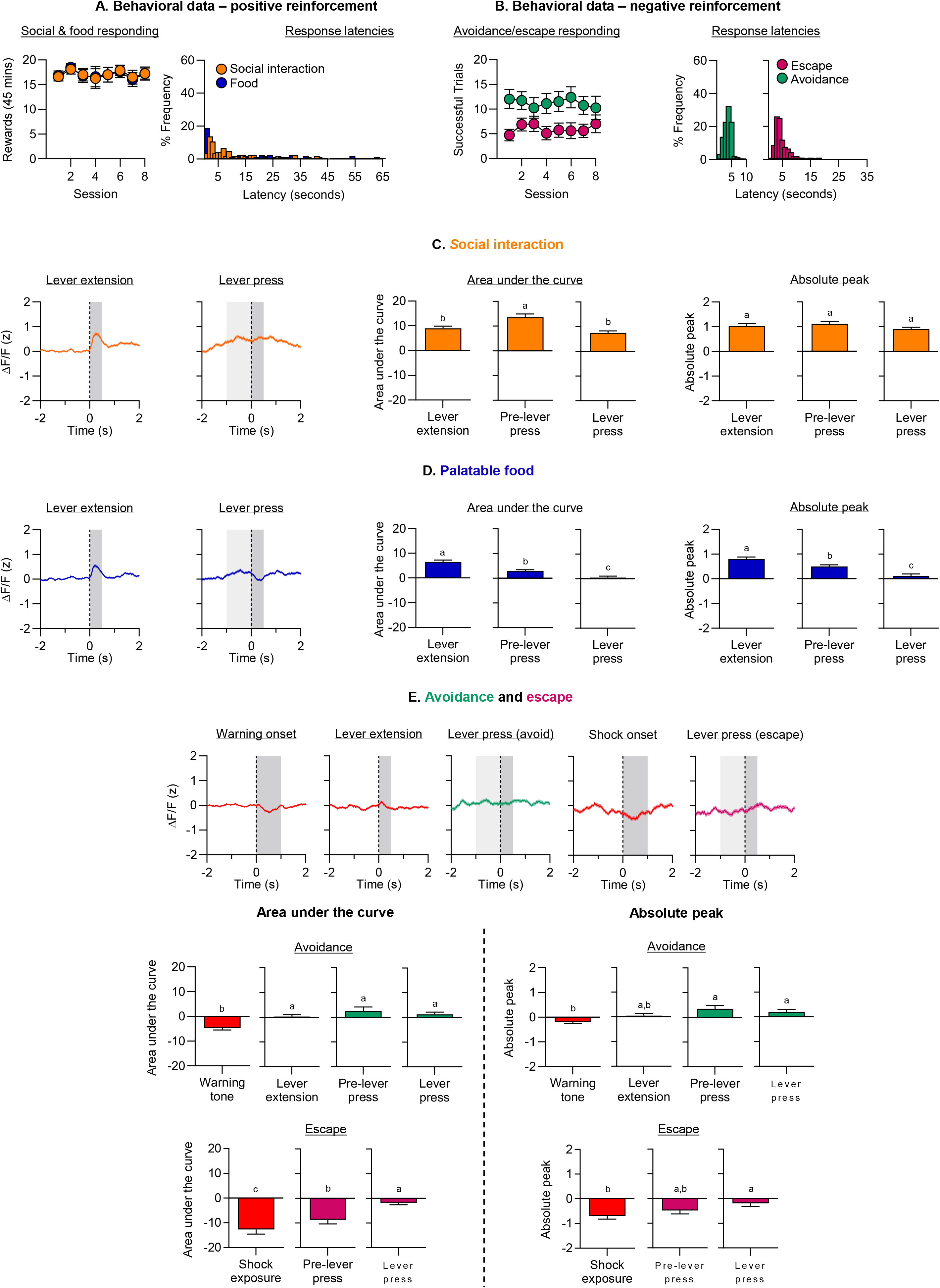
Dopamine activity in the NAc core during the three-reinforcer sessions. **(A)** Number of food and social interaction rewards earned and latency to lever-press following lever extension during three-reinforcer sessions. **(B)** Number of successful avoidance and escape trials and latency to lever-press following lever extension during three-reinforcer sessions. **(C)** Dopamine responses during social interaction trials, from lever extension through the lever-press (left), with corresponding AUC (middle) and absolute peak amplitude (right) measures for lever extension, the pre-press period, and the lever-press. **(D)** Dopamine responses during food trials, from lever extension through the lever-press (left), with corresponding AUC (middle) and absolute peak amplitude (right) measures for lever extension, the pre-press period, and the lever-press. **(E)** Dopamine responses during shock trials at, from left to right, warning-signal onset, lever extension, the avoidance response, shock onset, and the escape response. Letters indicate results of post hoc comparisons, ranked from highest (a) to lowest; those sharing at least one letter are not significantly different. n = 8 (4 females).

#### Fiber photometry recording data

For all photometry recordings, we determined the area under the curve (AUC; net dopamine signal amplitude deviation from 0) and the absolute peak amplitude (highest or lowest dopamine signal amplitude deviating from 0) at lever extension (to 0.5 s after), the period right before the lever-press (from 1 s before to event), and immediately following the lever-press (to 0.5 s after). We chose a 1-s epoch prior to lever-press to analyze based on our prior study (Chow et al., 2024b) (Note: using a 0.5 s does not change the overall results). For shock avoidance and escape trials, we additionally analyzed dopamine activity during the warning signal and from shock onset to 1-s after shock onset, respectively.

We first analyzed AUC and absolute peak responses separately for each reinforcer and session type (single-reinforcer vs. three-reinforcer sessions), with Event (nominal) as the within-subjects factor. For food and social interaction, Event included: lever insertion, the pre-lever-press period, and the post-lever-press period. For shock avoidance, Event included warning-signal onset, lever insertion, the pre-lever-press avoidance-response period, and the post-lever-press avoidance-response period. For shock escape, Event included shock onset, the pre-lever-press escape-response period, and the post-lever-press escape-response period.

We next analyzed these data with Event and Session Type (nominal: single-reinforcer vs. three-reinforcer sessions) as within-subjects factors and conducted post-hoc tests comparing corresponding events between session types.

Finally, we compared activity during lever insertion, the pre-lever press period, and the post-lever press period across the different reinforcers used, with Reinforcer Type (nominal: social interaction, food, shock avoidance, or shock escape) and Session Type (single-reinforcer vs. three-reinforcer sessions) as within-subjects factors.

### Single-reinforcer recordings

#### Social interaction

NAc core dopamine recordings during social self-administration showed a phasic increase in dopamine activity following lever extension, a gradual increase leading up to the lever-press, and a decrease toward baseline following the lever-press (Figure 2C). Dopamine levels were similar following lever extension and during the period preceding the lever-press.

#### Area under the curve (AUC)

The analysis showed a significant effect of Event [F(2, 239.1) = 10.3, p < 0.0001]. Post-hoc tests showed that AUC was higher following lever extension and during the pre-lever-press period than during the post-lever-press period.

#### Absolute peak

The analysis showed a significant effect of Event [F(2, 255.5) = 9.4, p = 0.0001]. Post-hoc tests showed that absolute peak responses were higher following lever extension and during the pre-lever press period than during the post-lever press period.

#### Palatable food

NAc core dopamine recordings during food self-administration showed a phasic increase in dopamine activity following lever extension, a further increase leading up to the lever-press, and a decrease following the lever-press (Figure 2D).

#### Area under the curve

The analysis showed a significant effect of Event [F(2, 210.7) = 131.6, p < 0.0001]. Post-hoc tests showed that AUC was highest following lever extension, followed by the pre-lever-press period and then the post-lever-press period.

#### Absolute peak

The analysis showed a significant effect of Event [F(2, 224.8) = 92.7, p < 0.0001]. Post-hoc tests showed the same pattern as the AUC analysis.

#### Shock avoidance

NAc core dopamine recordings during shock avoidance showed a phasic decrease in dopamine activity at warning-signal onset, followed by a similar decrease following lever extension. Dopamine activity remained relatively stable during the periods preceding and following the avoidance response (Figure 3C).

#### Area under the curve

The analysis showed a significant effect of Event [F(3, 645.5) = 60.2, p < 0.0001]. Post-hoc tests showed that AUC was lowest at warning-signal onset, followed by lever extension, and was higher during the pre- and post-avoidance-response periods.

#### Absolute peak

The analysis showed a significant effect of Event [F(3, 632.8) = 30.1, p < 0.0001]. Post hoc-tests showed that absolute peak responses were lowest at warning-signal onset and following lever extension.

#### Shock escape

NAc core dopamine recordings during shock escape showed a phasic decrease in dopamine activity at shock onset. Dopamine activity remained relatively stable during the periods preceding and following the escape response (Figure 3C).

#### Area under the curve

The analysis showed a significant effect of Event [F(2, 259.2) = 65.0, p < 0.0001]. Post-hoc tests showed that AUC was lowest at shock onset, followed by the pre-escape-response period, and was highest during the post-escape-response period.

#### Absolute peak

The analysis showed a significant effect of Event [F(2, 286.3) = 39.3, p < 0.0001]. Post-hoc tests showed that absolute peak responses were lowest at shock onset, whereas responses during the pre- and post-escape-response periods were similar.

### Three-reinforcer recordings

#### Social interaction

NAc core dopamine recordings during social self-administration showed a phasic increase in dopamine activity following lever extension and a gradual increase leading up to and through the lever-press (Figure 4C).

#### Area under the curve

The analysis showed a significant effect of Event [F(2, 249.5) = 12.0, p < 0.0001]. Post-hoc tests showed that AUC was higher during the pre-lever-press period than following lever extension or during the post-lever-press period.

#### Absolute peak

The analysis showed no significant effect of Event.

#### Palatable food

NAc core dopamine recordings during food self-administration showed a phasic increase in dopamine activity following lever extension, a further increase leading up to the lever-press, and a gradual decrease following the lever-press (Figure 4D).

#### Area under the curve

The analysis showed a significant effect of Event [F(2, 283.6) = 31.2, p < 0.0001]. Post hoc tests showed that AUC was highest following lever extension, followed by the pre-lever-press and post-lever-press periods.

#### Absolute peak

The analysis showed a significant effect of Event [F(2, 287.6) = 22.4, p < 0.0001]. Post-hoc tests showed the same pattern as the AUC analysis.

#### Shock avoidance

NAc core dopamine recordings during shock avoidance showed a phasic decrease in dopamine activity at warning-signal onset. Dopamine activity during the periods preceding and following the avoidance response was similar (Figure 4E).

#### Area under the curve

The analysis showed a significant effect of Event [F(3, 288.9) = 13.3, p < 0.0001]. Post-hoc tests showed that AUC was lowest at warning-signal onset.

#### Absolute peak

The analysis showed a significant effect of Event [F(3, 317.2) = 6.9, p < 0.0001]. Post-hoc tests showed that absolute peak responses were lowest at warning-signal onset. The response following lever extension was similar to responses at warning-signal onset and during the pre- and post-avoidance-response periods.

#### Shock escape

NAc core dopamine recordings during shock escape showed a phasic decrease in dopamine activity at shock onset. Dopamine activity during the periods preceding and following the escape response was similar (Figure 4E).

#### Area under the curve

The analysis showed a significant effect of Event [F(2, 114.3) = 28.6, p < 0.0001]. Post-hoc tests showed that AUC was lowest at shock onset, followed by the pre-escape-response period, and was highest during the post-escape-response period.

#### Absolute peak

The analysis showed a significant effect of Event [F(2, 128) = 5.7, p < 0.0001]. Post-hoc tests showed that absolute peak responses were lowest at shock onset. Responses during the pre-escape-response period were similar to those at shock onset and during the post-escape-response period.

### Single-reinforcer versus three-reinforcer recordings

#### Social interaction

NAc core dopamine recordings during social self-administration showed similar patterns and magnitudes of activity following lever extension and during the pre-lever-press period in single-and three-reinforcer sessions. However, dopamine responses diverged following the lever-press: dopamine levels decreased during the post-lever-press period in single-reinforcer sessions but remained elevated in three-reinforcer sessions (Figure 5A).

**Figure 5.**
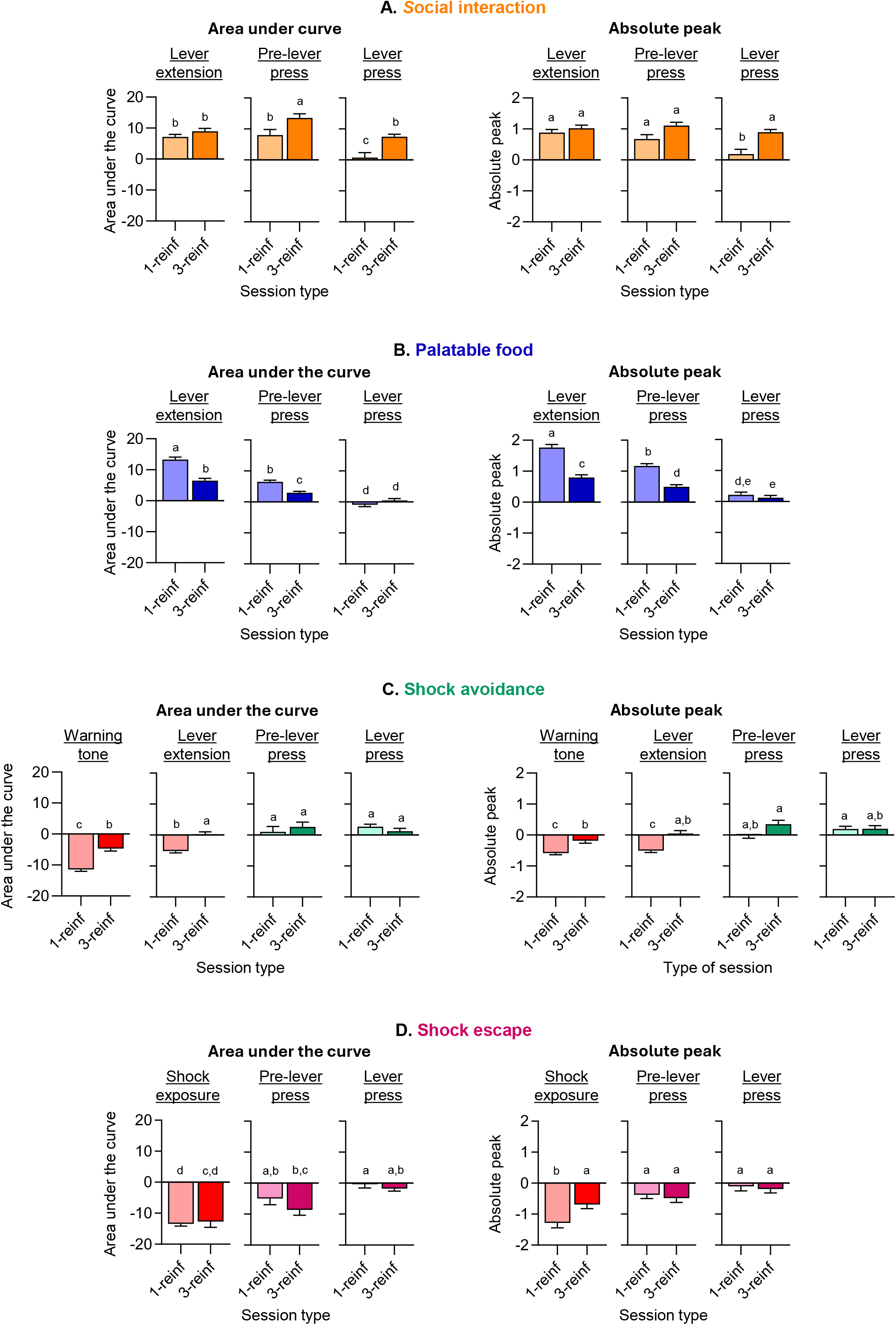
Comparison of dopamine responses between one-reinforcer and three-reinforcer sessions. Area under the curve and absolute peak amplitude for social interaction **(A)**, palatable food **(B)**, shock avoidance **(C)**, and shock escape **(D)**. Letters indicate results of post hoc comparisons, ranked from highest (a) to lowest; those sharing at least one letter are not significantly different. n = 8 (4 females).

#### Area under the curve

The analysis showed significant effects of Event [F(2, 251.4) = 19.2, p < 0.0001], Session Type [F(1, 139.2) = 15.0, p = 0.0002], and Event × Session Type interaction [F(2, 252.9) = 3.5, p = 0.032]. Post-hoc tests showed that the interaction was driven by the post-lever-press response, which decreased during single-reinforcer sessions but remained elevated during three-reinforcer sessions.

#### Absolute peak

The analysis showed significant effects of Event [F(2, 270.5) = 10.8, p < 0.0001], Session Type [F(1, 141.3) = 13.9, p = 0.0003], and Event × Session Type interaction [F(2, 277.6) = 4.1, p = 0.018]. Post-hoc tests showed the same pattern as the AUC analysis.

#### Palatable food

NAc core dopamine recordings during food self-administration showed similar patterns of dopamine activity following lever extension, during the pre-lever-press period, and during the post-lever press period in single- and three-reinforcer sessions (Figure 5B). However, responses following lever extension and during the pre-lever press period were greater during single-reinforcer sessions than during three-reinforcer sessions, whereas responses during the post-lever-press period did not differ between the session types.

#### Area under the curve

The analysis showed significant effects of Event [F(2, 260.1) = 138.9, p < 0.0001], Session Type [F(1, 122.9) = 29.7, p < 0.0001], and Event × Session Type interaction [F(2, 240.9) = 30.50, p < 0.0001]. Post-hoc tests showed that the interaction was driven by progressive decreases in the dopamine response from lever extension through the lever-press. The post-lever-press response did not differ between single- and three-reinforcer sessions.

#### Absolute peak

The analysis showed significant effects of Event [F(2, 267.7) = 100.5, p < 0.0001], Session Type [F(1, 125.8) = 58.1, p < 0.0001], and Event × Session Type interaction [F(2, 246.2) = 20.2, p < 0.0001]. Post-hoc tests showed the same pattern as the AUC analysis.

#### Shock avoidance

NAc core dopamine recordings during shock avoidance showed decreases in dopamine activity at warning-signal onset and following lever extension in both single- and three-reinforcer sessions (Figure 5C). Responses at these events differed between Session Types, whereas responses during the pre- and post-avoidance-response periods were similar across Session Types.

#### Area under the curve

The analysis showed significant effects of Event [F(3, 581.3) = 25.5, p < 0.0001], Session Type [F(1, 170.2) = 25.3, p < 0.0001], and Event × Session Type interaction [F(3, 569.4) = 3.5, p = 0.016]. Post-hoc tests showed that responses at warning-signal onset and following lever extension were lower during single-reinforcer sessions, whereas responses during the pre- and post-avoidance-response periods did not differ between session types.

#### Absolute peak

The analysis showed significant effects of Event [F(3, 633.4) = 48.6, p < 0.0001], Session Type [F(1, 206.3) = 15.7, p = 0.0001], and Event × Session Type interaction [F(3, 625.6) = 6.9, p = 0.0002]. Post-hoc tests showed a similar pattern to the AUC analysis.

#### Shock escape

NAc core dopamine recordings during shock escape showed a decrease in dopamine activity at shock onset in both single- and three-reinforcer sessions (Figure 5D). Responses at shock onset differed between Session Types, whereas responses during the pre- and post-escape-response periods were similar across Session Types.

#### Area under the curve

The analysis showed significant effects of Event [F(2, 45.0) = 51.2, p < 0.0001] and Event × Session Type interaction [F(2, 104.5) = 4.5, p = 0.014]. Post-hoc tests showed that AUC was lower at shock onset than during the pre- and post-escape-response periods. In addition, the response at shock onset was lower during single-reinforcer sessions than during three-reinforcer sessions.

#### Absolute peak

The analysis showed significant effects of Event [F(2, 146) = 21.1, p < 0.0001] and the Event × Session Type interaction [F(2, 121.1) = 4.7, p = 0.011]. Post-hoc tests showed that the absolute peak response at shock onset during single-reinforcer sessions was lower than responses during all other shock-escape-related events.

#### Comparison of dopamine responses during operant responding for positive and negative reinforcers

We next compared dopamine responses across the different reinforcer outcomes during lever extension, pre-lever-press, and lever-press. For each event, we analyzed AUC and absolute peak data using Reinforcer Outcome (nominal) and Session Type (nominal; one-reinforcer, three-reinforcers) as within-subjects factors. For lever extension, the reinforcer outcomes were social interaction, food, and shock. For pre-lever-press and lever-press, the reinforcer outcomes were social interaction, food, shock avoidance, and shock escape.

#### Lever extension

NAc core dopamine responses following lever extension differed across reinforcer outcomes and session types. During single-reinforcer sessions, the food-associated lever elicited the largest increase in dopamine activity, whereas the shock-avoidance-associated lever elicited the largest decrease. During three-reinforcer sessions, the food- and social-interaction-associated levers elicited similar increases in dopamine activity, and both responses were greater than the response to the shock-avoidance-associated lever (Figure 6A,B, left).

**Figure 6.**
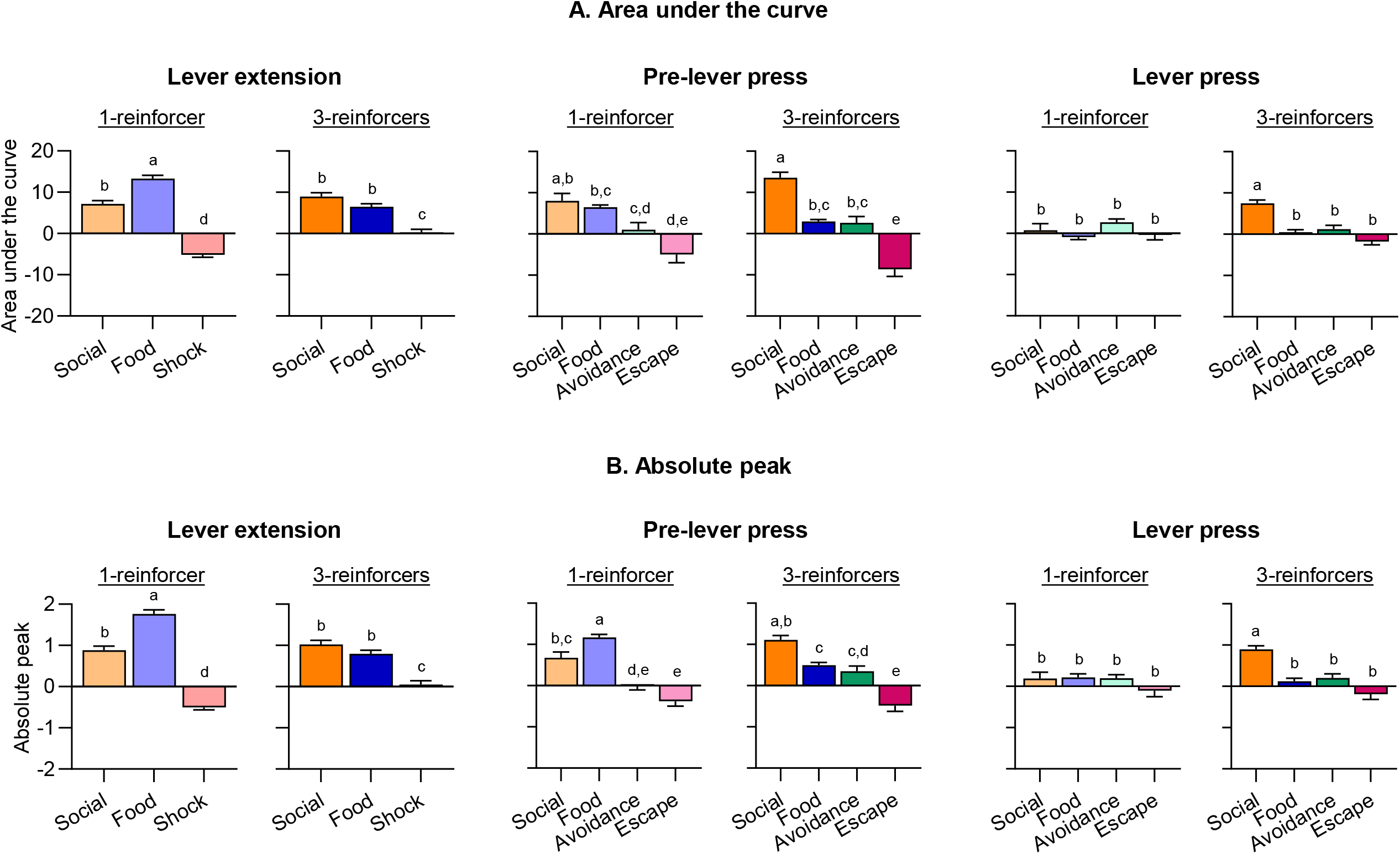
Comparison of dopamine responses across reinforcement conditions and recording events during single-reinforcer and three-reinforcer sessions. Area under the curve **(A)** and absolute peak amplitude **(B)** during lever extension, the pre-lever-press period, and the post-lever-press period for social interaction, palatable food, shock avoidance, and shock escape. Letters indicate results of post hoc comparisons, ranked from highest (a) to lowest; those sharing at least one letter are not significantly different. n = 8 (4 females).

#### Area under the curve

The analysis showed significant effects of Reinforcer Outcome [F(2, 187.8) = 185.2, p < 0.0001] and Reinforcer Outcome × Session Type interaction [F(2, 191.6) = 40.6, p < 0.0001]. Post-hoc tests showed that during single-reinforcer sessions, the food-associated cue elicited a larger phasic dopamine response than the social-interaction and shock-avoidance cues, and the social-interaction cue elicited a larger response than the shock-avoidance cue. During three-reinforcer sessions, the food- and social-interaction cues elicited comparable phasic dopamine responses, and both elicited larger responses than the shock-avoidance cue.

#### Absolute peak

The analysis showed significant effects of Reinforcer Outcome [F(2, 285.7) = 189.6, p < 0.0001] and Reinforcer Outcome × Session Type interaction [F(2, 283.5) = 43.0, p < 0.0001]. Post-hoc-tests showed the same pattern as the AUC analysis.

Together, these findings show that a cue (lever extension) signaling positive reinforcers elicited larger phasic dopamine responses than the cue signaling shock avoidance. The food-associated cue elicited the largest dopamine response during single-reinforcer sessions but not during three-reinforcer sessions.

#### Pre-lever-press

Dopamine responses during the pre-lever-press period differed across the different reinforcer outcomes and session types. Overall, responses were greatest for positive reinforcers and lowest for shock escape, with shock avoidance producing intermediate responses (Figure 6A,B, middle).

#### Area under the curve

The analysis showed significant effects of Reinforcer Outcome [F(3, 396) = 37.5, p < 0.0001] and Reinforcer Outcome × Session Type interaction [F(3, 417.8) = 3.9, p = 0.0090]. Post-hoc tests showed that the AUC was highest for social interaction and progressively decreased across the other reinforcer outcomes, with the lowest response observed for shock escape.

#### Absolute peak

The analysis showed significant effects of Reinforcer Outcome [F(3, 371.5) = 54.4, p < 0.0001] and Reinforcer Outcome × Session Type interaction [F(3, 397.9) = 11.6, p < 0.0001]. Post-hoc tests showed that the absolute peak was highest for food during single-reinforcer sessions and lowest for shock escape under both Session Types. The overall pattern was food during single-reinforcer sessions > social interaction and food during three-reinforcer sessions > shock avoidance > shock escape.

#### Post-lever-press

Dopamine responses during the post-lever-press period were generally similar across the reinforcer outcomes and session types, except for the elevated response following social interaction during three-reinforcer sessions (Figure 6A,B, right).

#### Area under the curve

The analysis showed significant effects of Reinforcer Outcome [F(3, 399.3) = 11.3, p < 0.0001] and Reinforcer Outcome × Session Type interaction [F(3, 435.1) = 7.8, p < 0.0001]. Post-hoc tests showed that the AUC following the lever-press for social interaction during three-reinforcer sessions was higher than the AUC for all other outcomes.

#### Absolute peak

The analysis showed significant effects of Reinforcer Outcome [F(3, 375) = 10.8, p < 0.0001] and Reinforcer Outcome × Session Type interaction [F(3, 407.1) = 5.9, p = 0.0006]. Post-hoc tests showed the same pattern as the AUC analysis.

## Discussion

We examined dopamine activity in the NAc core of rats that learned to lever-press for positive reinforcement (social interaction with a peer and high-carbohydrate palatable food) and negative reinforcement (to avoid or escape mild footshock). Several main findings emerged:

First, during social self-administration, dopamine activity showed phasic increases following lever insertion and gradual increases preceding lever pressing. These responses were modestly greater when all three reinforcers were available than when social interaction was the only available reinforcer (Figure 5A). These findings replicate and extend our previous results obtained using a social self-administration procedure (Chow et al., 2024b).

Second, palatable food self-administration produced a similar pattern of dopamine activity. However, dopamine responses were approximately twofold greater when food was the only available reinforcer than when all three reinforcers were available during the periods following lever insertion and preceding lever-pressing, but not at the time of lever-pressing itself (Figure 5B). In sessions in which only a single reinforcer was available, the dopamine response to lever insertion was significantly greater for palatable food than for social interaction. In contrast, during sessions in which all three reinforcers were available, dopamine responses during the pre-lever-press period and at lever-pressing were greater for social interaction than for palatable food (Figure 6).

Third, during shock avoidance/escape, dopamine activity showed phasic decreases following warning onset, lever insertion, and shock onset. These decreases were also greater when shock avoidance/escape was the only available reinforcer than when all three reinforcers were available (Figure 5C, D).

Together, these findings indicate that NAc core dopamine signaling differentiates positive from negative reinforcement and that the availability of other reinforcers modulates dopamine responses in a reinforcer-specific manner.

### Nucleus accumbens dopamine response to cues associated with operant reinforcers

Our finding that lever insertion, a cue predicting the availability of either social interaction or palatable food, increases NAc core dopamine transmission is consistent with previous studies. These studies used microdialysis, fiber photometry, or fast-scan cyclic voltammetry during operant procedures involving addictive drugs (Kiyatkin and Stein, 1996; Tran-Nguyen et al., 1998; Phillips et al., 2003; Stuber et al., 2005), palatable food (Roitman et al., 2004; Saddoris et al., 2015; Mohebi et al., 2019), or social interaction (Chow et al., 2024b) (for review, see Wise (2004)).

A consistent finding in our study was a phasic decrease in NAc core dopamine in response to cues predicting the negative reinforcer (the warning tone and lever insertion). The cue-related dopamine decreases contrast with previous reports of increased phasic NAc dopamine responses to cues predicting shock avoidance (Oleson et al., 2012; Gentry et al., 2016; Wenzel et al., 2018). Wenzel et al. (2018) also showed that stimulation of VTA dopamine neurons projecting to NAc core enhanced avoidance responding. Similarly, a microdialysis study using 1-min sampling intervals reported increased NAc dopamine in response to a shock-paired cue (Young, 2004). Finally, a non-operant avoidance study also reported increased phasic NAc core dopamine responses to shock-paired cues (Lopez et al., 2025).

Thus, in studies using different recording methods and behavioral procedures, shock-predictive cues have generally been associated with increased NAc dopamine signaling, making the phasic dopamine decreases observed in our study particularly notable. The discrepant responses to shock-predictive cues may reflect procedural differences. Unlike previous studies, which examined operant avoidance/escape in isolation, our rats were trained to respond for two positive reinforcers and one negative reinforcer. Additionally, the shock used in our study was milder (0.14-0.28 mA for 0.5 s) than that used in previous studies (0.42-0.56 mA) (Oleson et al., 2012; Gentry et al., 2016; Wenzel et al., 2018). These differences in reinforcement context and shock intensity may contribute to the distinct NAc dopamine responses observed across studies.

Phasic dopamine signals in the NAc have been proposed to encode aspects of reward value (Schultz et al., 2015), and, more broadly, NAc dopamine has been proposed to encode the incentive-motivational effects of reward-predictive cues (Stewart et al., 1984; Berridge and Robinson, 2003). From this perspective, our finding that lever insertion elicited a larger dopamine response when it predicted palatable food than when it predicted social interaction during the single-reinforcer sessions is consistent with the greater reinforcing efficacy of the food reward under our experimental conditions. In previous studies, the palatable food used here (Calu et al., 2014) was strongly preferred over social interaction; additionally, rats worked harder to obtain this food reinforcer, as indicated by their willingness to complete higher fixed-ratio requirements, providing an additional measure of its greater reinforcing efficacy (Chow et al., 2022).

However, the relationship between the phasic dopamine response to a reward-predictive cue and the relative reinforcing efficacy of the predicted reward was not straightforward. If the larger dopamine response to the food-predictive cue during the single-reinforcer sessions reflected the greater reinforcing efficacy or incentive motivational value of food, a similar difference might have been expected when all three reinforcers were available. This was not observed. Thus, although the dopamine response to the food-predictive cue was larger than the response to the social-interaction cue under the single-reinforcer condition, this difference was not maintained when other reinforcers were also available. These findings suggest that NAc core phasic dopamine responses to positive-reinforcer-predictive cues are not determined solely by the relative reinforcing efficacy of the reinforcer.

In contrast, the qualitative difference between the dopamine responses to cues associated with positive and negative reinforcement was highly consistent across conditions. Specifically, the warning tone and lever insertion associated with shock avoidance/escape elicited dopamine decreases, whereas lever insertion associated with social interaction and palatable food elicited dopamine increases, regardless of whether the relevant reinforcer was presented alone or together with the other reinforcers. These findings suggest that phasic NAc core dopamine signaling can reliably discriminate between cues predicting positive versus negative operant outcomes, at least under our experimental conditions.

### Methodological considerations

Several methodological considerations should be considered in interpreting our data. First, the one-reinforcer and three-reinforcer recording sessions were not counterbalanced. Therefore, the conclusion that the availability of other reinforcers modulates dopamine responses to a given reinforcer should be interpreted cautiously, because differences between the two session types could reflect an order effect. We consider this possibility relatively unlikely, however, because the effects differed across reinforcers: the availability of other reinforcers was associated with lower dopamine responses during palatable food or shock avoidance/escape sessions but higher dopamine responses during social-interaction sessions.

Second, direct comparisons of dopamine responses to different behavioral events across reinforcers require caution. For food and social interaction, lever insertion was the primary predictive cue. In contrast, during shock avoidance/escape, lever insertion was preceded by a warning tone that itself elicited a decrease in dopamine activity. Thus, the dopamine response to lever insertion during shock avoidance/escape cannot be directly compared with the response to lever insertion during food or social-interaction sessions.

Third, comparisons of dopamine responses during the pre-lever-press period and at lever-pressing are complicated by differences in the latency between lever insertion and lever-press. The latency was short for palatable food (median, ∼1 s during recording) but longer and more variable for social interaction and shock avoidance/escape (see figures). Therefore, the temporal windows used to characterize these responses may not represent equivalent behavioral periods across reinforcers.

Fourth, fiber photometry provides a population-level measure of dopamine dynamics with limited spatial and cellular resolution. Consequently, it remains unknown whether the recorded signals reflect activity in common or distinct neuronal populations that contribute to responding to reinforcer-associated cues.

Fifth, dopamine responses surrounding lever insertion and pressing in operant tasks may reflect a combination of motivational, sensory, cognitive, and motor processes. Therefore, the present measurements cannot be attributed exclusively to reinforcement-related processes. This issue is particularly relevant because the three reinforcers differed in their sensory and behavioral properties.

Finally, the small sample size does not allow sex-related differences in dopamine signaling under the present experimental conditions.

## Conclusions

Our findings suggest that phasic NAc core dopamine signaling differentiates cues predicting positive versus negative reinforcement. In contrast, its ability to distinguish between different positive reinforcers according to their relative reinforcing efficacy appears to be more limited and dependent on the broader reinforcement context. Thus, although NAc core phasic dopamine signaling may reflect aspects of reward value and incentive motivation, it is not a simple readout of the relative reinforcing efficacy of positive reinforcers.

## Notes

Funding and Disclosure: This research was supported by the Intramural Research Program of NIDA (ZIA-DA000434-25, YS) and support from grants (K99DA062788, JJC). The authors declare that they do not have any conflicts of interest (financial or otherwise). The contributions of the NIH author(s) are considered Works of the United States Government. The findings and conclusions presented in this paper are those of the author(s) and do not necessarily reflect the views of the NIH or the U.S. Department of Health and Human Services.

### Competing Interest Statement

The authors have declared no competing interest.

